# Iterative Spatial Resolution Enhancement in Imaging Mass Spectrometry via Hydrogel Tissue Expansion and Multimodal Image Fusion

**DOI:** 10.64898/2026.06.03.729902

**Authors:** Elijah D. Mayo, Jacob M. Samuel, Yingchan Guo, Audrey Ciccone, Zhongling Liang, Boone M. Prentice

**Affiliations:** Department of Chemistry, University of Florida, Gainesville, FL. 32608

**Author notes:** Address correspondence to: Dr. Boone M. Prentice, 214 Leigh Hall, PO Box 117200, Department of Chemistry University of Florida Gainesville, FL 32611, USA.

## Abstract

The pixel size of imaging mass spectrometry (IMS) is fundamentally limited by several factors, including the diameter of the incident probe and the raster step size of the sample stage. We have previously demonstrated that hydrogel-based tissue expansion, originally developed for microscopy (ExM), can also be adapted for imaging mass spectrometry to physically magnify the size of the tissue. Expansion imaging mass spectrometry (ExIMS) uses a superabsorbent hydrogel to isotropically expand thin tissue sections, which can then be sampled via imaging mass spectrometry, resulting in improved effective spatial resolution. Separately, multimodal image fusion has been used to computationally upsample the effective spatial resolution in imaging mass spectrometry by predictively mapping mass spectrometric intensity values to the smaller diameter pixel sizes of a microscopy image of the same tissue section. Here, we present ExFusion, a unified workflow that combines these two approaches by computationally fusing structurally detailed fluorescent ExM and chemically detailed lipid ExIMS data obtained from the same 9.4-fold expanded mouse brain tissue. Following a 10-fold upsampling from image fusion, multimodal expansion image fusion enabled prediction of MS images at a ∼106 nm pixel size on a commercial mass spectrometer using a 10 μm raster step size. At this resolution, lipids in the Purkinje cells of the cerebellum are clearly defined with intracellular distributions.

## INTRODUCTION

Imaging mass spectrometry (IMS) provides a powerful, label-free technique for spatial molecular profiling of biological tissues and cell cultures.^1^ Unlike immunofluorescence microscopy, which relies on antibody labeling, IMS enables the untargeted analysis of metabolites, lipids, and peptides with specificity determined by ionization efficiency and mass resolution rather than antibody affinity.^2^ Still, imaging MS has limitations. For example, pixel size is limited by factors such as the diameter of the incident ionization probe and the stage raster step. State-of-the-art commercial matrix-assisted laser/desorption ionization (MALDI) imaging MS instruments are limited to ∼5 μm pixel size.^3-5^ This limits the ability of imaging MS to provide cellular-level detail, especially when compared to immunofluorescence or hematoxylin and eosin (H&E) microscopy. While instrument modifications can be made to decrease sampling raster width to ∼1 μm, these modifications are usually expensive and invasive.^6-9^ Therefore, developing accessible, cost-effective approaches to high spatial resolution imaging MS would represent a significant contribution to the spatial biology community.

Hydrogel tissue expansion can be implemented in imaging MS (i.e., ExIMS) to physically magnify the tissue, thereby improving effective spatial resolution in proportion to the linear expansion factor.^10-16^ Initially validated in microscopy (ExM), tissue expansion isotropically enlarges tissues by crosslinking the monomers of a superabsorbent hydrogel to the lysine residues of transmembrane and structural proteins.^17-20^ When submerged in water, the expansion of the gel decrowds cellular structures, physically magnifying the gel so that it can be imaged at a higher effective spatial resolution using the same instrument hardware (*i*.*e*., same stage raster step and same laser diameter). We have recently shown that lipids, even without a lipid-specific covalent anchor to the hydrogel network, are largely retained when expanded ∼4-fold.^15^ Other recent reports have shown that tissues can be expanded up to ∼10-fold without signal loss in MALDI and DESI imaging MS.^21, 22^ Expansion imaging MS has also been extended to the high spatial resolution mapping of metabolites, peptides, and N-glycans.^14, 16, 23, 24^ In microscopy, workflows achieving greater than 10-fold expansion have been developed,^25, 26^ but this may reduce sensitivity and induce analyte delocalization in ExIMS.^15^ Given these limitations, expansion alone may be insufficient to achieve subcellular resolution in ExIMS. Pairing tissue expansion with complementary resolution enhancement strategies may therefore be a promising path toward achieving nanoscale pixel sizes.

Multimodal image fusion has been used to computationally upsample the digital pixel size in imaging mass spectrometry by predictively mapping imaging MS intensity values to the features of a higher spatial resolution image, most commonly microscopy.^27, 28^ We have previously presented a workflow that uses supervised and deep learning models to perform image fusion using MALDI imaging MS and H&E microscopy images acquired from the same tissue section.^29^ After accurate image alignment, a cross-modality relationship between the imaging MS and H&E images is mathematically modeled using a linear regression, partial-least squares (PLS) regression, random forest regression, or 2D convolutional neural network. From this computational image fusion, a predicted image of the mass spectrometry ion intensities is created at the digital pixel size of the H&E image. This enables better imaging MS resolution of tissue morphology that could not have been determined in the original MS image.

Here, we introduce a workflow that combines tissue expansion with multimodal image fusion (ExFusion). First, using hydrogel tissue expansion, we physically magnified the tissue to reduce the effective sampling pixel size. Next, using multimodal image fusion, we computationally refined the digital pixel size based on a fluorescent microscopy image of the same tissue area. By sequentially applying physical resolution enhancement by ExIMS, then computational resolution enhancement image fusion, ExFusion enables an iterative improvement to the overall spatial resolution for imaging mass spectrometry, without any alterations to the commercially available mass spectrometer used to perform imaging MS.

## METHODS

The ExIMS and image fusion workflows used in this study are based on the methods we have previously reported for each, as well as the anchoring and gelation protocol reported by Klimas et al. to achieve the almost 10-fold expansion factor.^15, 20, 29^ The ExFusion workflow consists of three stages: tissue preparation and expansion for the physical sampling pixel size enhancement, multimodal imaging using fluorescence microscopy and MALDI IMS, and computational image fusion for the digital pixel size enhancement. The staining for fluorescence is performed after the initial gelation, but before the full expansion in water, as phosphate buffered saline causes some shrinking to the expanded gel, which risks lipid delocalization and stain degradation. After full expansion, the gel must be dried completely before entering the vacuum environment of the mass spectrometer. Imaging the gel after drying is not required for microscopy, but aids in the registration process.

**Figure 1.**
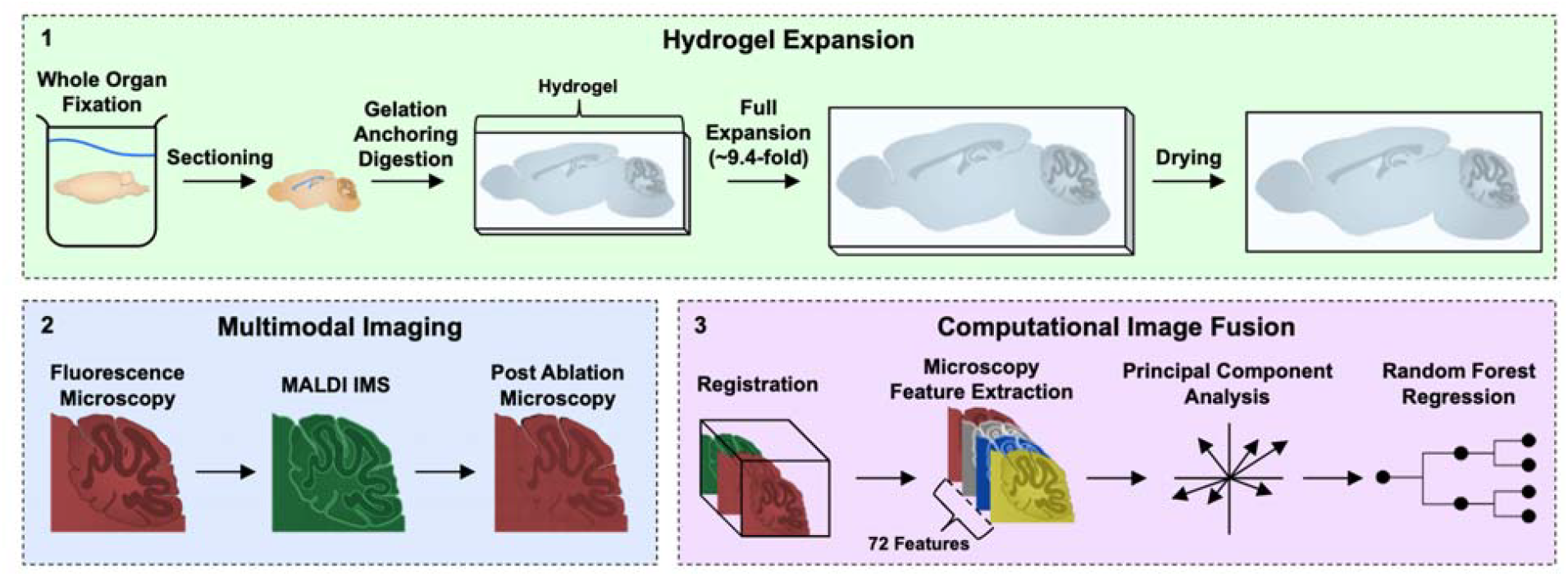
ExFusion workflow consists of three main steps: (1) isotropic enlargement of tissue via hydrogel expansion, (2) multimodal imaging, and (3) computational image sharpening via image fusion. In Step 1, a whole mouse brain is fixed and cryoprotected, cryosectioned, embedded in a hydrogel network, homogenized via proteolytic digestion, stained, expanded in water, and dried using a combination of vacuum desiccation and nitrogen gas. In Step 2, the expanded mouse brain section is imaged using fluorescence microscopy prior to imaging mass spectrometry. After MALDI imaging mass spectrometry, an additional fluorescence image is acquired to visualize laser ablation marks. In Step 3, the MS image is first aligned to the post-MS ablation image, and then to the pre-MS fluorescence image. From the fluorescence microscopy image, a range of image features are extracted and dimensionally reduced through principal component analysis. A cross-modality relationship is then modeled using a random forest regression between the principal components derived from the fluorescence image and the ion intensity distributions derived from the MS image. This mathematical relationship then enables prediction of the original MS image to a higher-spatial resolution MS image, effectively reducing the pixel size and sharpening the image. Created with BioDraws.com.

### Chemicals and Reagents

HPLC-grade water, phosphate buffered saline (PBS), sucrose, sodium chloride, Atto 633 NHS Ester, 4′,6-diamidino-2-phenylindole (DAPI), and solvents for matrix spraying were obtained from ThermoFisher Scientific (Waltham, MA, USA). Methacrolein, 1,5-diaminonaphthalene (DAN), 4-hydroxy-2,2,6,6-tetramethylpiperidin-1-oxyl (4HT), ammonium persulfate (APS), acrylamide, tetramethylenediamine (TEMED), and tris-hydrochloride were purchased from Sigma-Aldrich (St. Louis, MO, USA). Proteinase K was purchased from New England Biosciences (Ipswich, MA, USA). Mouse brain (C57BL/6) tissue was acquired from BioIVT (Westbury, NY, USA). Paraformaldehyde was acquired from Boster Bio (Pleasanton, CA, USA). Sodium acrylate was purchased from Santa Cruz Biotechnology (Dallas, TX, USA).

### Tissue Preparation

Mouse brain tissue was fixed in a 4% paraformaldehyde and 0.1% glutaraldehyde solution in 1x PBS overnight at 4°C, cryoprotected by 30% (w/v) sucrose in 1× PBS at 4°C until the brain sank, then frozen with isopentane chilled with dry ice, and stored at 80°C. Tissue sectioning was performed on a Leica CM3050S cryomicrotome (Leica Biosystems, Wetzlar, Germany). Fixed mouse brain was sectioned transversely to 30 μm thickness in pairs and placed in a well of 1x PBS, then mounted onto a microscope slide for ExM/ExIMS or onto an indium-tin oxide (ITO) slide for normal fluorescence staining and imaging mass spectrometry analysis.

### Gelation

Tissue sections were incubated for five minutes at 4°C with 90 μL of a dual anchoring and monomer solution (1x PBS, 1% (w/v) NaCl, 34% (w/v) sodium acrylate, 10 % (w/v) acrylamide, 0.01 % (w/w) MBAA) with 0.2 % (w/v), 4HT, 2% (w/v) APS, 2% (w/v) TEMED, and 0.1 % methacrolein) added to the solution immediately before gelation. While still immersed in the monomer solution, an additional 90 μL of the monomer solution was placed on the tissue sections, and they were placed in a custom-built gelation chamber and allowed to polymerize at 4°C for 48 hours. The gelation chamber was then removed, and the embedded tissues were digested with 4 U/mL proteinase K in pH 8 digestion buffer (50 mM Tris-HCl and 0.8 M NaCl) overnight at room temperature.

### Microscopy

After removing the digestion buffer, tissues were rinsed once and washed with 1x PBS for 10 minutes. Primary amines were labeled with 5 µg/mL NHS ester ATTO633 in 1x PBS for 30 minutes at room temperature. Tissues were then rinsed once and washed with 1x PBS for 10 minutes. Nuclear DNA was stained with 0.5 µg/mL DAPI in 1x PBS for 5 minutes at room temperature, followed by a final wash with 1x PBS, three times for 10 minutes each. Tissues were then fully expanded by submersion in excess d.d. H_2_O, desiccated at 1.3 Torr until corners of the gel were flattened to the slide, and then left to dry under nitrogen gas flow overnight. Post drying, tissues were imaged with a 10x objective using an Axio Imager M2 microscope (Carl Zeiss, Jenna, Germany) with DAPI and 633 nm channels. Post-MS, laser ablation patterns were imaged for registration using the same channels. Both pre-MS fluorescence and post-MS ablation images were downsampled to 100-fold the pixel density of the imaging MS images and exported as TIFF images using the ZEN software (Carl Zeiss, Jenna, Germany) image processing feature.

### Imaging Mass Spectrometry

A 1,5-diaminonaphthalene (DAN) matrix layer was applied using an M5 sprayer (HTX Technologies, LLC, Chapel Hill, NC) at 80°C nozzle temperature and four passes. All lipid imaging mass spectrometry experiments were conducted on a timsTOF fleX mass spectrometer (Bruker Daltonics, Billerica, MA) equipped with a Smartbeam 3D Nd:YAG laser system (10 kHz, 355 nm). Positive ion mode lipid imaging was conducted using 25 laser shots at 30% power. Images were acquired at 10 µm raster width. Total ion count (TIC) normalization was performed, and the imaging dataset was exported as an .imzml file from SciLS Lab software (Bruker Daltonics, Billerica, MA). Lipid identification was performed by searching against the LIPID MAPS database. Individual ion images were exported from the .imzml file as singular .csv files using an in-house Python script.

### Image Fusion

Image registration was performed with an in-house code in MATLAB using the pre-MS fluorescence image to visualize tissue structures and post-MS image to visualize the laser ablation pattern.^29, 30^ The mass spectrometry image is first aligned to the ablation image by matching 3–4 pixels to the corresponding laser ablation marks in the matrix coating, which are clearly visualized by microscopy. Next, the post-MS image is registered to the pre-MS image by identifying 4-5 matching tissue structure points. This sequential alignment links the coordinate system of the MS image to that of the pre-MS fluorescence image. Non-affine transformations are applied throughout, as all images originate from the same tissue region.

Feature extraction and multimodal fusion were performed with an in-house Python script. A total of 72 features were extracted per pixel after feature initialization, local range filtering, and entropy filtering (**Supplemental Table S1**). Principal component analysis (PCA) was performed on these features to create a hierarchical list of features, and 55 principal components was found to be optimal for this workflow. Multimodal image fusion was performed on the MS image and the 55 principal components from the microscopy image using a random forest regression model (RandomForestRegressor function from the Scikit-learn library) with 200 decision trees and no maximum depth. Lipid ion distribution fidelity in the predicted image was assessed by comparison with the original imaging mass spectrometry data. Evaluation metrics included the average residuals and the Pearson correlation coefficient (PCC). Average residuals were calculated using **Equation 1**:

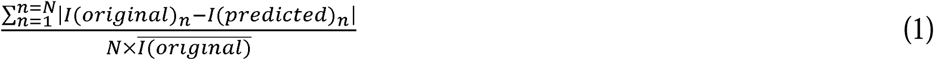

where N is the total number of pixels, I(original)n is the intensity value of the nth pixel in the original mass spectrometry image, and I(predicted)n is the intensity value of the nth pixel in the predicted mass spectrometry image. A residual image was also generated by plotting the per-pixel absolute difference between the original and predicted ion intensities, normalized by the average original ion intensity. The PCC was computed (Pearsonr function from SciPy) on the flattened imaging MS and predicted imaging MS matrices. Finally, prediction robustness was assessed by generating a per-pixel 95% confidence interval (CI) image derived from the distribution of predictions made by individual decision trees within the trained random forest model. To better match the residuals image, we chose to map model uncertainty at the 95% CI, rather than model certainty.

## RESULTS

### Spatial Resolution Enhancement

As a proof-of-concept, imaging mass spectrometry and fluorescence microscopy were performed on the same hydrogel-expanded mouse brain cerebellum tissue (**Figure 2**). Improvements in spatial resolution can be qualitatively assessed by monitoring the sharpness of small features in the tissue. For example, the Purkinje cell layer is a single-cell stratum in the brain, and these cells have diameters of 20-30 μm.^31^ While this cell layer can be probed using imaging mass spectrometry of the unexpanded tissue acquired using a 10 μm pixel size, the cells are not well resolved (**Figure 2A**). Image fusion alone substantially improves spatial sharpening, but the delineation of intracellular morphology is not achieved (**Figure 2C**). Tissue expansion results in a linear expansion factor of ∼9.4-fold compared to the original unexpanded tissue, as determined by averaging distances between identical biological landmarks in expanded versus unexpanded serial sections. Using a 10 μm stage step size, this results in an effective pixel size of ∼1.06 μm in the expanded tissue (**Figure 2D**). Performing computational image fusion on this expanded tissue results in a further 10-fold improvement in spatial resolution, resulting in a ∼94-fold improvement in pixel size. This allows for prediction of the mouse cerebellum image to a pixel size of ∼106 nm (**Figure 2F**). This pixel size is well below the accepted laser focusing and stage precision limits of this system (*i*.*e*., ∼5 µm).^32^ The 106 nm pixel size is also well below the limit needed for subcellular resolution. The combination of these two methods, hydrogel tissue expansion and multimodal image fusion, provides a superior spatial enhancement than either method has previously achieved alone.

**Figure 2.**
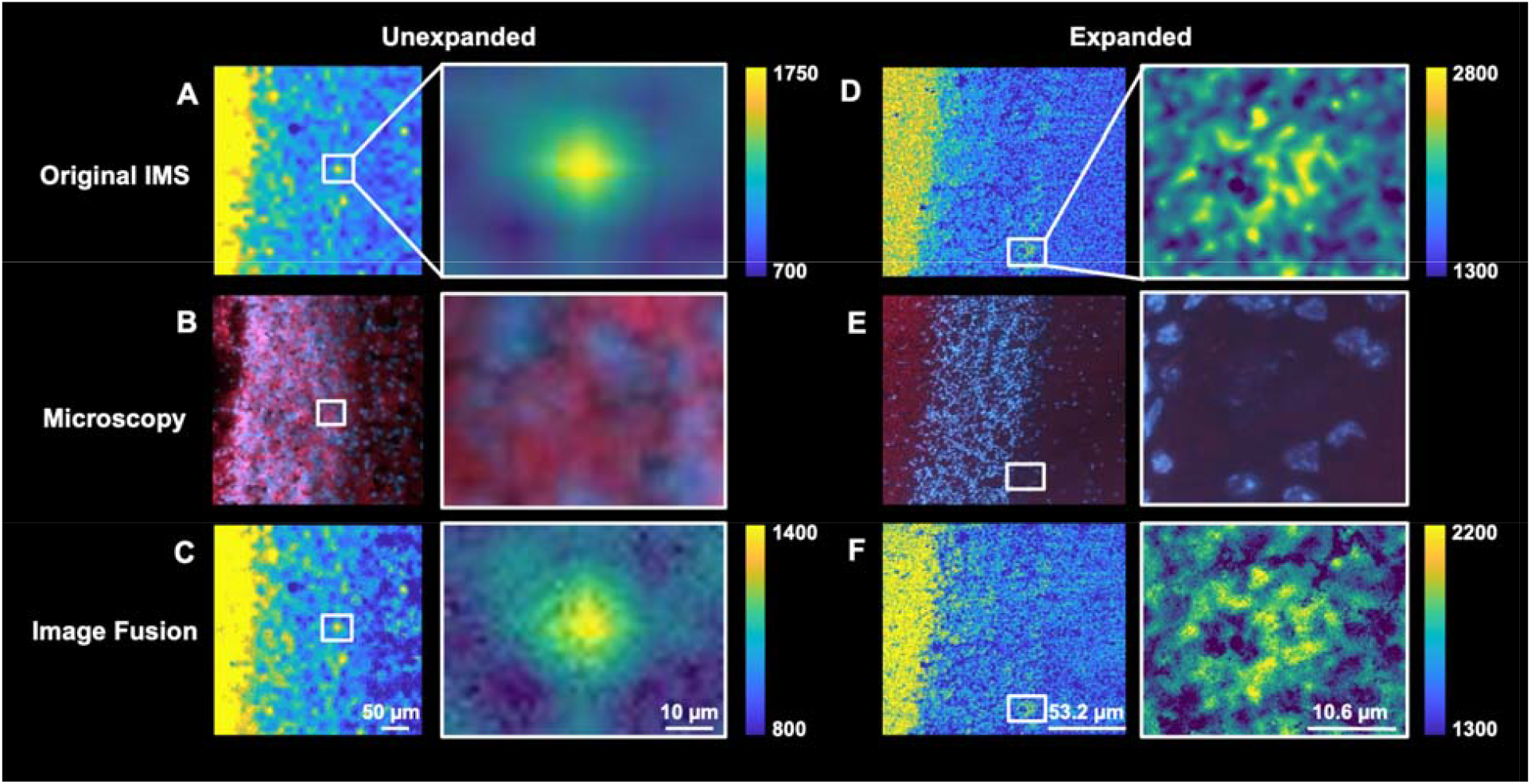
ExFusion enables subcellular spatial resolution: (A-C) Unexpanded and (D-F) ∼9.4-fold expanded mouse brain cerebellum analysis is performed using (A, D) imaging mass spectrometry, (B, E) fluorescence microscopy, and (C, F) image fusion for *m/z* 788.616, identified as [PC 36:1 + H]^+^ (-0.88 ppm). Images to the right in each panel are zoomed-in regions of the white box in the corresponding left image in each panel and identify regions representing a Purkinje cell. All MS images are acquired using a 10 μm stage step and are illustrated using bilinear interpolation (Python 3.11). Microscopy channels are DAPI (blue) for nucleic DNA and NHS Ester AF 633 (red) for general protein content. Scale bars are acquired from microscopy images (ZEN software, Carl Zeiss, Jenna, Germany) and divided by the measured expansion factor where appropriate.

The full resolving power of ExFusion is enabled by both the quantitative pixel size enhancement from each method (*i*.*e*., tissue expansion and image fusion) as well as the physical separation of overlapping structures afforded by expansion. When these techniques are combined to analyze the distribution of lipids at the intracellular level, the ∼106 nm enhanced pixel size and physical decrowding of cerebellum cell layers enables clear and consistent delineation of individual nuclei and soma boundaries (**Figure 3**). Importantly, neither tissue expansion nor image fusion enhancement causes significant distortions to the *in situ* lipid distributions. Saturated (**Figure 3A**), monounsaturated (**Figure 3B and 3C**), and polyunsaturated lipids (**Figure 3D**) are generally retained in their original cell layers across the cerebellum. The latter is noteworthy because polyunsaturated lipids have been reportedly more prone to delocalization during tissue expansion, but are well retained here within the Purkinje cell and molecular layers (**Figures 3D**).^15^ Similarly, image fusion retains intracellular distributions for both lowly (**Figure 3A and 3B**) and highly (**Figure 3C and 3D**) abundant lipids, suggesting that the prediction model is not biased towards the microscopy features used for training.

**Figure 3.**
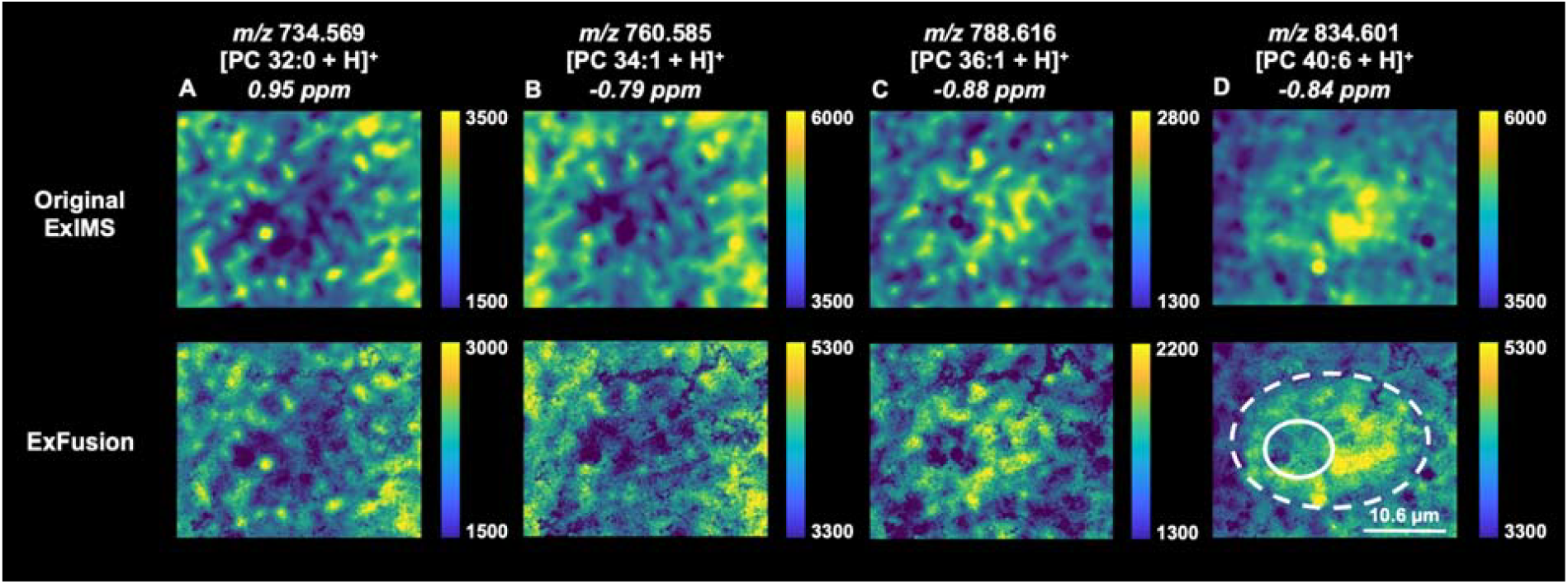
ExFusion reveals intracellular lipid distributions in Purkinje cells: ExFusion is demonstrated for (A) [PC 32:0 +H]^+^, (B) [PC 34:1 +H]^+^, (C) [PC 36:1 +H]^+^, and (D) [PC 40:6 +H]^+^. (Top) Imaging mass spectrometry analyses of ∼9.4-fold expanded mouse brain cerebellum tissue are acquired using a 10 μm stage step size. (Bottom) Multimodal image fusion results in further spatial resolution enhancement. The solid white circle outlines the approximate nucleus boundary and the dotted white circle outlines the approximate soma boundary.

### Image Fusion Performance

One significant modification in our multimodal imaging workflow compared to conventional image fusion workflows is the replacement of H&E staining and brightfield imaging with fluorescence staining and microscopy. This change was made to accommodate the hydrophilic environment of the expanded gel and remove the risk of washing away lipids during eosin staining. DAPI is used as a fluorescent replacement for hematoxylin and NHS Ester AF633 is used as a replacement for eosin in order to remain consistent with the morphology attained in conventional image fusion workflows. To verify this modification, and tissue expansion in general, had no significant effect on the image fusion model’s performance, we assessed the model’s prediction error, its ability to preserve correlation with the original imaging mass spectrometry data, and its prediction robustness. Prediction error was quantified by calculating both the mean residual across the image and by generating a pixel-by-pixel map of the residual error. Correlation was calculated using the Pearson correlation coefficient (PCC) between the original and predicted MS images. Prediction robustness was evaluated by generating a pixel-by-pixel map of the 95% confidence interval (CI) based on the model’s uncertainty. This CI map draws from the confidence interval image proposed by van de Plas, *et al*. and our previous standard deviation image, but here leverages the existing structure of the random forest without requiring retraining or additional sampling.^27, 29^ Together, these metrics offer a comprehensive view of model performance, providing interpretable, spatially resolved measures to assess fusion accuracy and robustness at the per-pixel level. For example, the abundance of [PC 36:1 + H]^+^ is highest in the white matter and Purkinje cells and lowest in the granular and molecular layers (**Figure 2, 3C, 4C**). This spatial distribution remains largely consistent in the cerebellum between the unexpanded and expanded tissues, both before and after image fusion. This observation is supported by the low mean residuals, high PCC scores, and comparable distributions of error and uncertainty across different lipid distributions (**Figure 4**). These evaluations indicate that the image fusion prediction is not overfitting to the image features captured in microscopy and is using well the fine morphological structures to upsample the digital pixel size. Additionally, the similarity in residual and uncertainty mapping images suggests that the model’s performance is only slightly impaired in tissue areas where the microscopy and imaging mass spectrometry data misalign (*e*.*g*., the tissue tear in the right side of the image field). This could be due to minor lipid delocalization during tissue expansion or due to regions of higher ion abundance, such as the white matter (**Figure 4B and 4C**) and Purkinje cell layer (**Figure 4C and 4D**). Overall, lipids in these cerebellum cell layers show relatively low error and high correlation between the original ExIMS and predicted images, indicating that lipids retain their spatial fidelity during tissue expansion and image fusion.

**Figure 4.**
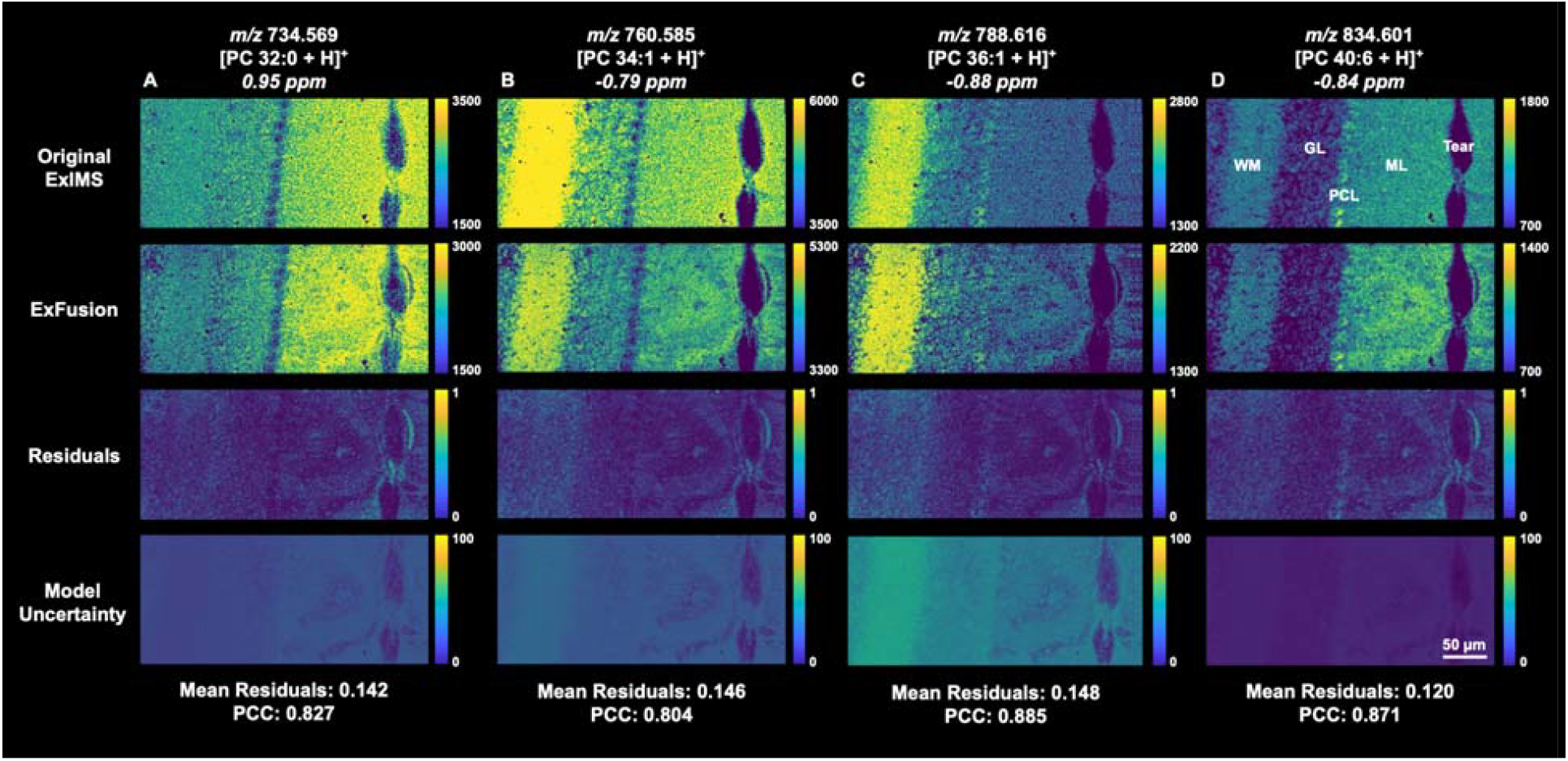
Image fusion performance remains consistent across cerebellum lipid distributions: ExFusion performance is evaluated for (A) [PC 32:0 +H]^+^, (B) [PC 34:1 +H]^+^, (C) [PC 36:1 +H]^+^, and (D) [PC 40:6 +H]^+^. (Top Row) ExIMS is first performed on a ∼9.4-fold expanded mouse brain cerebellum tissue are acquired using a 10 μm stage step size. (Second Row) Multimodal image fusion is then performed using a random forest model. ExFusion performance is assessed using pixel-by-pixel mapping of (Third Row) the residuals and (Bottom Row) the 95% CI. Labeled tissue regions are ML = Molecular Layer, GL = Granular Layer, PCL = Purkinje Cell Layer, WM = White Matter.

## CONCLUSIONS

By integrating hydrogel-based tissue expansion with multimodal image fusion, ExFusion offers a novel and effective combination for iteratively improving the spatial resolution of imaging mass spectrometry experiments. This sequential application of physical and computational resolution enhancement enables a level of spatial detail that surpasses the capabilities of either method alone. The application of ExFusion to mouse brain tissue enabled a 94-fold improvement in pixel size without requiring changes to the commercial instrument (*i*.*e*., the MALDI laser or sampling stage), and enabled intracellular imaging below the diffraction limit of the MALDI laser system. The enhanced spatial resolution provided by ExFusion enables clear visualization of fine anatomical features, such as intracellular Purkinje cell components, which are poorly resolved in conventional imaging mass spectrometry. While we achieved a pixel size of ∼106 nm, further improvements in spatial resolution are attainable through the use of higher expansion factor gels or larger differences in imaging MS and microscopy pixel size, but these methods require careful evaluation of lipid distribution fidelity at each step. Given the robustness of image fusion using fluorescence microscopy, improved visualization may be possible using better staining protocols for subcellular structures, though these may induce lipid delocalization if permeabilization is required. The accuracy of image fusion after tissue expansion was validated here by calculating the average residuals and PCC metrics as well as by mapping the residuals and 95% confidence interval uncertainty. These criteria demonstrate that ExFusion does not significantly alter image fusion accuracy. However, the results of any image fusion prediction should be used cautiously, especially in areas of higher error and model uncertainty. Overall, the spatial resolution enhancement achieved by ExFusion offers promise in other high spatial resolution applications, including single-cell segmentation, clustering, and intracellular biomolecule co-localization.

## ACKNOWLEDGEMENTS

This work was supported by the Alfred P. Sloan Foundation and the Eastman Foundation.

## REFERENCES

(1) Norris, J. L.; Caprioli, R. M. Analysis of tissue specimens by matrix-assisted laser desorption/ionization imaging mass spectrometry in biological and clinical research. Chem Rev 2013, 113 (4), 2309–2342. DOI: 10.1021/cr3004295.

(2) Zaima, N.; Hayasaka, T.; Goto-Inoue, N.; Setou, M. Matrix-assisted laser desorption/ionization imaging mass spectrometry. Int. J. Mol. Sci. 2010, 11 (12), 5040–5055. DOI: 10.3390/ijms11125040.

(3) Zhang, H.; Lu, K. H.; Ebbini, M.; Huang, P.; Lu, H.; Li, L. Mass spectrometry imaging for spatially resolved multi-omics molecular mapping. Npj Imaging 2024, 2 (1), 20. DOI: 10.1038/s44303-024-00025-3.

(4) Feenstra, A. D.; Dueñas, M. E.; Lee, Y. J. Five Micron High Resolution MALDI Mass Spectrometry Imaging with Simple, Interchangeable, Multi-Resolution Optical System. J Am Soc Mass Spectrom 2017, 28 (3), 434–442. DOI: 10.1007/s13361-016-1577-8.

(5) Spraggins, J. M.; Djambazova, K. V.; Rivera, E. S.; Migas, L. G.; Neumann, E. K.; Fuetterer, A.; Suetering, J.; Goedecke, N.; Ly, A.; Van de Plas, R.; et al. High-performance molecular imaging with MALDI trapped ion-mobility time-of-flight (timsTOF) mass spectrometry. Anal. Chem. 2019, 91 (22), 14552–14560. DOI: 10.1021/acs.analchem.9b03612 From NLM.

(6) Spengler, B.; Hubert, M. Scanning microprobe matrix-assisted laser desorption ionization (SMALDI) mass spectrometry: instrumentation for sub-micrometer resolved LDI and MALDI surface analysis. J Am Soc Mass Spectrom 2002, 13 (6), 735–748. DOI: 10.1016/S1044-0305(02)00376-8.

(7) Zavalin, A.; Yang, J.; Hayden, K.; Vestal, M.; Caprioli, R. M. Tissue protein imaging at 1 μm laser spot diameter for high spatial resolution and high imaging speed using transmission geometry MALDI TOF MS. Anal. Bioanal. Chem. 2015, 407, 2337–2342. DOI: 10.1007/s00216-015-8532-6

(8) Spivey, E. C.; McMillen, J. C.; Ryan, D. J.; Spraggins, J. M.; Caprioli, R. M. Combining MALDI-2 and transmission geometry laser optics to achieve high sensitivity for ultra-high spatial resolution surface analysis. J Mass Spectrom 2019, 54 (4), 366–370. DOI: 10.1002/jms.4335.

(9) Niehaus, M.; Soltwisch, J.; Belov, M. E.; Dreisewerd, K. Transmission-mode MALDI-2 mass spectrometry imaging of cells and tissues at subcellular resolution. Nat Methods 2019, 16 (9), 925–931. DOI: 10.1038/s41592-019-0536-2.

(10) Bai, Y.; Zhu, B.; Oliveria, J. P.; Cannon, B. J.; Feyaerts, D.; Bosse, M.; Vijayaragavan, K.; Greenwald, N. F.; Phillips, D.; Schürch, C. M.; et al. Expanded vacuum-stable gels for multiplexed high-resolution spatial histopathology. Nat Commun 2023, 14 (1), 4013. DOI: 10.1038/s41467-023-39616-w.

(11) Hung, Y. L. W.; Xie, C.; Wang, J.; Diao, X.; Li, R.; Wang, X.; Qiu, S.; Fang, J.; Cai, Z. Expansion Strategy-Driven Micron-Level Resolution Mass Spectrometry Imaging of Lipids in Mouse Brain Tissue. CCS Chem 2024, 6, 2662–2670. DOI: 10.31635/ccschem.024.202404002

(12) Chen, L. C.; Lee, C.; Hsu, C. C. Towards developing a matrix-assisted laser desorption/ionization mass spectrometry imaging (MALDI MSI) compatible tissue expansion protocol. Anal. Chim. Acta 2024, 1297, 342345. DOI: 10.1016/j.aca.2024.342345.

(13) Chan, Y. H.; Pathmasiri, K. C.; Pierre-Jacques, D.; Hibbard, M. C.; Tao, N.; Fischer, J. L.; Yang, E.; Cologna, S. M.; Gao, R. Gel-assisted mass spectrometry imaging enables sub-micrometer spatial lipidomics. Nat Commun 2024, 15 (1), 5036. DOI: 10.1038/s41467-024-49384-w.

(14) Zhang, H.; Ding, L.; Hu, A.; Shi, X.; Huang, P.; Lu, H.; Tillberg, P. W.; Wang, M. C.; Li, L. TEMI: tissue-expansion mass-spectrometry imaging. Nat Methods 2025, 22 (5), 1051–1058. DOI: 10.1038/s41592-025-02664-9.

(15) Samuel, J. M.; Baby, N. M.; Mayo, E. D.; Yan, T.; Liang, Z.; Prentice, B. M. Examination of lipid distributions in hydrogel-expanded mouse brain tissue using imaging mass spectrometry. Anal Chim Acta 2025, 1377, 344629. DOI: 10.1016/j.aca.2025.344629.

(16) Wang, F.; Sun, C.; Wu, T. W.; Fu, Y.; Fan, Y.; Zhao, S.; Huang, K.; Pan, Z.; Lu, Y.; Han, J. R.; et al. iPEX enables micrometre-resolution deep spatial proteomics via tissue expansion. Nature 2026, 649 (8096), 505–514. DOI: 10.1038/s41586-025-09734-0.

(17) Chen, F.; Tillberg, P. W.; Boyden, E. S. Optical imaging. Expansion microscopy. Science 2015, 347 (6221), 543–548. DOI: 10.1126/science.1260088.

(18) Tillberg, P. W.; Chen, F.; Piatkevich, K. D.; Zhao, Y.; Yu, C. C.; English, B. P.; Gao, L.; Martorell, A.; Suk, H. J.; Yoshida, F.; et al. Protein-retention expansion microscopy of cells and tissues labeled using standard fluorescent proteins and antibodies. Nat. Biotechnol. 2016, 34 (9), 987–992. DOI: 10.1038/nbt.3625.

(19) Min, K.; Cho, I.; Choi, M.; Chang, J. B. Multiplexed expansion microscopy of the brain through fluorophore screening. Methods 2020, 174, 3–10. DOI: 10.1016/j.ymeth.2019.07.017 From NLM.

(20) Klimas, A.; Gallagher, B. R.; Wijesekara, P.; Fekir, S.; DiBernardo, E. F.; Cheng, Z.; Stolz, D. B.; Cambi, F.; Watkins, S. C.; Brody, S. L.; et al. Magnify is a universal molecular anchoring strategy for expansion microscopy. Nat. Biotechnol. 2023, 41 (6), 858–869. DOI: 10.1038/s41587-022-01546-1.

(21) Xie, C.; Wang, J.; Diao, X.; Guo, L.; Lam, T. K.-Y.; Li, R.; Chen, Y.; Zhang, Y.; Wang, X.; Fang, J.; et al. Ten-Fold Expansion MALDI Mass Spectrometry Imaging of Tissues and Cells at 500 Nm Resolution. bioRxiv 2024. DOI: 10.1101/2024.10.20.619316.

(22) Xie, C.; Wang, J.; Diao, X.; Wang, X.; Cai, Z. Single-Cell Resolution DESI Mass Spectrometry Imaging through 10-Fold Sample Expansion. bioRxiv 2024. DOI: 10.1101/2024.10.21.619369

(23) Zimmerman, T. A.; Rubakhin, S. S.; Sweedler, J. V. MALDI Mass Spectrometry Imaging of Neuronal Cell Cultures. J. Am. Soc. Mass Spectrom. 2011, 22 (5), 828–836. DOI: 10.1007/s13361-011-0111-2.

(24) Tucker, K. R.; Serebryannyy, L. A.; Zimmerman, T. A.; Rubakhin, S. S.; Sweedler, J. V. The modified-bead stretched sample method: Development and application to MALDI-MS imaging of protein localization in the spinal cord. Chemical Science 2011, 2 (4), 785–795. DOI: 10.1039/c0sc00563k.

(25) Chang, J. B.; Chen, F.; Yoon, Y. G.; Jung, E. E.; Babcock, H.; Kang, J. S.; Asano, S.; Suk, H. J.; Pak, N.; Tillberg, P. W.; et al. Iterative expansion microscopy. Nat. Methods 2017, 14 (6), 593–599. DOI: 10.1038/nmeth.4261.

(26) Wang, S.; Shin, T. W.; Yoder, H. B.; McMillan, R. B.; Su, H.; Liu, Y.; Zhang, C.; Leung, K. S.; Yin, P.; Kiessling, L. L.; et al. Single-shot 20-fold expansion microscopy. Nat. Methods 2024, 21 (11), 2128–2134. DOI: 10.1038/s41592-024-02454-9.

(27) Van de Plas, R.; Yang, J.; Spraggins, J.; Caprioli, R. M. Image fusion of mass spectrometry and microscopy: a multimodality paradigm for molecular tissue mapping. Nat. Methods 2015, 12 (4), 366–372. DOI: 10.1038/nmeth.3296.

(28) Guo, L.; Zhu, J.; Wang, K.; Cheng, K. K.; Xu, J.; Dong, L.; Xu, X.; Chen, C.; Shah, M.; Peng, Z.; et al. Multimodal Image Fusion Offers Better Spatial Resolution for Mass Spectrometry Imaging. Anal. Chem. 2023, 95 (25), 9714–9721. DOI: 10.1021/acs.analchem.3c02002.

(29) Liang, Z.; Guo, Z.; Sharma, A.; McCurdy, C. R.; Prentice, B. M. Multimodal Image Fusion Workflow Incorporating MALDI Imaging Mass Spectrometry and Microscopy for the Study of Small Pharmaceutical Compounds. Anal. Chem. 2024, 96 (29), 11869–11880.

(30) Patterson, N. H.; Tuck, M.; Van de Plas, R.; Caprioli, R. M. Advanced registration and analysis of MALDI imaging mass spectrometry measurements through autofluorescence microscopy. Anal. Chem. 2018, 90 (21), 12395–12403. DOI: 10.1021/acs.analchem.8b02884 From NLM.

(31) Hammond, C.; Monique, E. The chemical synapses. In Cellular and Molecular Neurophysiology, 4 Ed.; Academic Press, 2015.

(32) Esselman, A. B.; Patterson, N. H.; Migas, L. G.; Dufresne, M.; Djambazova, K. V.; Colley, M. E.; Van de Plas, R.; Spraggins, J. M. Microscopy-Directed Imaging Mass Spectrometry for Rapid High Spatial Resolution Molecular Imaging of Glomeruli. J. Am. Soc. Mass Spectrom. 2023, 34 (7), 1305–1314. DOI: 10.1021/jasms.3c00033.

