## Supplemental Information for "Iterative Spatial Resolution Enhancement in Imaging Mass Spectrometry via Hydrogel Tissue Expansion and Multimodal Image Fusion"

**Affiliations:**

#Address correspondence to:

Dr. Boone M. Prentice

214 Leigh Hall

PO Box 117200

Department of Chemistry

University of Florida

Gainesville, FL 32611, USA

| **Feature Number** | **Feature/Feature Generating Function** |
| --- | --- |
| 0-2  3-5  6-8  9-11  12-14  15  16  17  18  19  20  21  22  23  24-47  48-71 | RGB  Lab  HSV  NTSC  YCbCr  Grayscale  Laplacian of Gaussian  Local Std Dev  Gradient Maximum  Squared x-Gradient  Squared y-Gradient  xy-Gradient product  RGB Channel Maximum  Canny Edge Detection  Local Range Filter  Entropy Range Filter |

**Supplemental Table 1**. Per-pixel features extracted from the microscopy image for image fusion. Principal component analysis was performed on these extracted features.
